# Sponges, ctenophores and the statistical significance of syntenies

**DOI:** 10.1101/2025.05.07.652388

**Authors:** Richard R. Copley

**Affiliations:** Laboratoire de Biologie du Développement, de Villefranche-sur-mer (LBDV), Sorbonne Université, CNRS UMR7009, 06230 Villefranche-sur-mer, France

## Abstract

Shared fusions between ancestral chromosomal linkage groups have previously been used to support phylogenetic groupings, notably sponges with cnidarians and bilaterians to the exclusion of ctenophores, rendering ctenophores the sister group to all other animals. The linkage groups used to identify these fusions were assessed for statistical significance relative to a model of randomly shuffled genes. I argue that the method of random shuffling treated all species as equally distant from each other, and so over-estimated the significance of the observed linkages. I calculate alternative statistics, and further argue that there are likely to be real linkage groups which are not identified as significant. If linkage groups are not supported statistically, they cannot reliably be used to identify shared derived chromosomal rearrangements, and hence phylogenetic hypotheses derived from them are suspect.

## 1 Introduction

Whether the ctenophores or sponges are sister group to the other animals has been the subject of continuing dispute since the first ‘phylogenomic’ era study to include ctenophores [1]. In that work, Dunn and colleagues assembled a multigene, multispecies sequence alignment and found several surprising phylogenetic relationships, among which, ctenophores as the earliest branching animal phylum attracted much interest. Since then, there has been some degree of back and forth between camps purporting to show ctenophores or sponges in this position [2, 3, 4, 5, 6, 7, 8, 9, 10]. Very broadly speaking, analyses focussed on complex phylogenetic models have tended to show sponges sister, whereas those focussed on assembling large data sets, ctenophores (but see [11]). Other sources of data besides sequence alignments have been explored (e.g. gene content [12, 13]), but these results have not had wide influence. Gene synteny data have recently been proposed as a possible independent source of informative phylogenetic markers. Schultz and co-workers [14] (henceforth SSR23) used inferred Ancestral Linkage Groups (ALGs) to hypothesise a number of polarised chromosome fusion events that were uniquely shared between sponges and the clade of cnidarians and bilaterians (i.e. parahoxozons), thus supporting ctenophores sister to other animals. SSR23 tested the statistical significance of the initial inferred ALGs through random permutation of chromosomal gene order. I suggest that the procedure they adopted leads to inflated estimates of significance, and that when more appropriate tests are used, support for the conservation of some ALGs disappears or is sensitive to the use of closely related genomes or the method of orthology identification.

SSR23 defined synteny groups with respect to four species quartets: a unicellular outgroup, a sponge, a ctenophore and a cnidarian or bilaterian. The number of orthologs, *N*, shared by potential synteny groups was assessed for significance by randomly permuting the gene order across all chromosomes within every species of the quartet, and counting the number of times a group of *N* or- thologs was observed. These values were used as estimates of the False Discovery Rate (FDR). There are two problems with this procedure. Firstly, by permuting all species, it takes no account of the fact that the species show greatly differing levels of phylogenetic relatedness. To see that this is a problem, imagine the hypothetical case of adding another species whose synteny is perfectly conserved with one already in the dataset, as if adding an extra wheel to a fruit machine. To correctly simulate the biological reality, these wheels should spin, or chromosomes be permuted, in lockstep. Instead, if all chromosomes are randomly permuted, the statistical significance of any synteny group in the unpermuted data will be lowered (i.e. made more significant) by more random failures, despite the underlying evolutionary history remaining the same.

The second problem with the procedure of SSR23 is the synthesis of the permuted data into a set of FDRs for a given gene group size (e.g. five orthologs have an FDR of *x*). These values give a probability for a group size averaged across all chromosomes. We might reasonably expect, however, that the statistical significance should depend on the number of shared genes relative to the sizes of the chromosomes that the linkages are observed on. A group of five genes shared on small chromosomes of the four species is less likely than the same group size shared by the largest chromosomes.

While the first of these problems, that of permutation of all species, will lead to more erroneously significant results, the effect of the second is less clear *a priori* as it will depend upon which chromosomes are likely syntenic. To investigate this further, I implemented an alternative approach that addresses both issues.

## 2 Results

### 2.1 Shuffling all species inflates statistical significance

In order to illustrate the effects of different permutation strategies ^1^ I developed code to calculate the number of times a particular combination of chromosomes was supported by shared orthologs. Using the data of SSR23, this exactly reproduced the sizes of the SS23 shared ortholog groups. I then made a dataset of orthologs with chromosomal locations for the cnidarian *Nematostella vectensis* and the sponge *Corticium candelabrum*, adding a fake third species by copying the *Nematostella* gene locations to a set of new chromosome names and made-up gene ids, which had a one-to-one mapping to the *Nematostella* data. To calculate the random distribution of chromosome combinations, I repeatedly shuffled the gene to chromosome mapping for a given species. I either permuted just the *Corticium* gene locations (i.e. the outgroup) relative to the two identical species, or the locations of all species (Figure 1a). When all species are shuffled, the frequency with which random chromosomal groups are supported by larger numbers of genes is greatly reduced, which would make observed ‘real’ groupings more significant. When the data of SSR23 [14] are used for the quartet of *Capsaspora owczarzaki, Hormiphora californiensis, Ephydatia muelleri and Rhopilema esculentum*, shuffling all species gives numbers comparable to those found in SSR23: groups of 5 linked orthologs are seen extremely rarely and groups of 8 never seen. In contrast, shuffling just the *C*.*owc* ortholog locations yields larger linked groups (Figure 1b).

**Figure 1:**
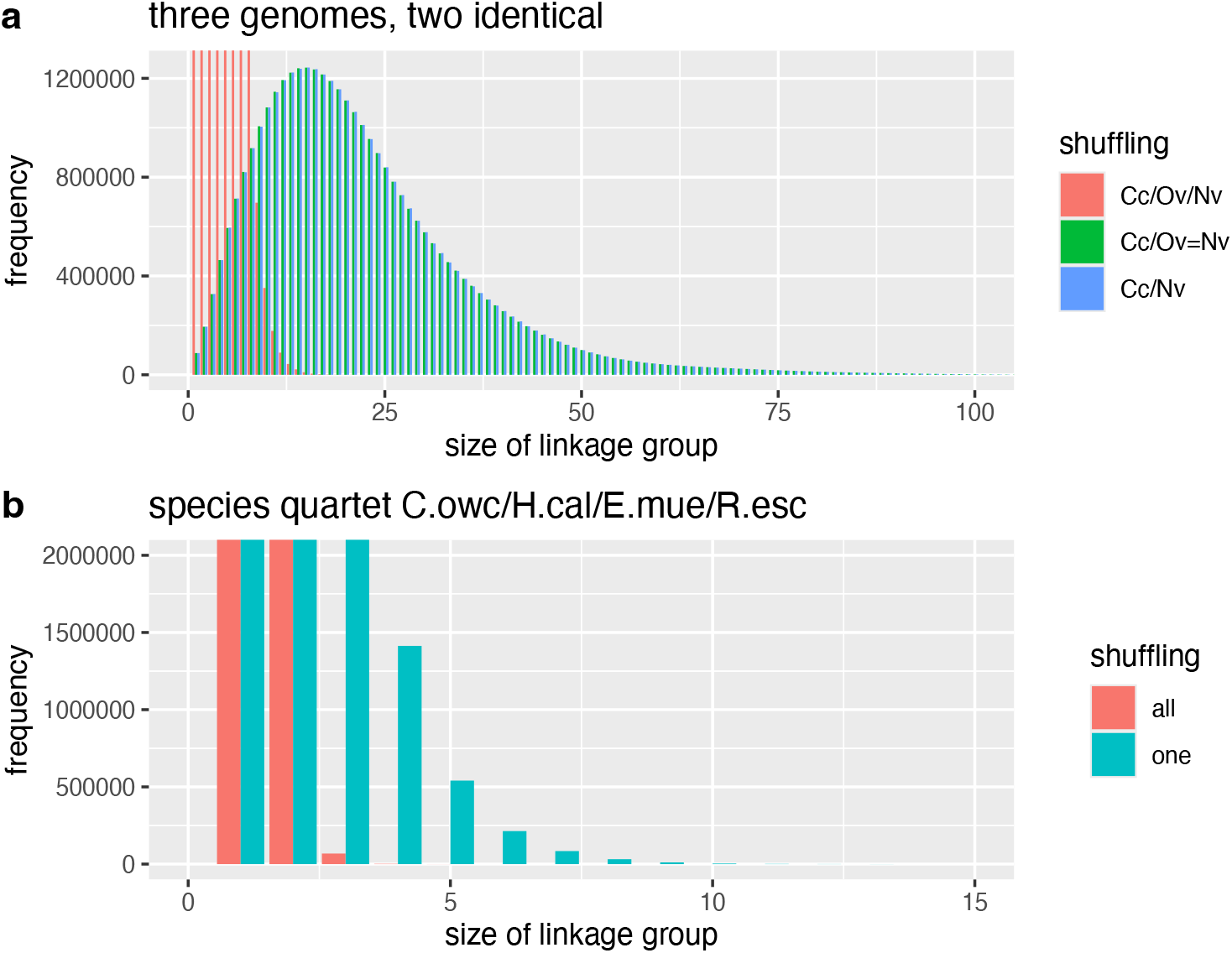
Effects of shuffling just one species *vs*. all species. The barplot bins show the frequency of a given number of linked orthologs for 100000 sets of randomized gene location data. In **a** the blue bars illustrate a pair of species, with one shuffled (*C*.*can/N*.*vec*). The green and red bars a trio of species where two are identical (*C*.*can/O*.*vec=N*.*vec*). In the green series, only the locations of *C*.*can* genes were shuffled. In the red series the locations of all genes were shuffled. The desired biological behaviour, an identical distribution to a pair of species, is seen with the shuffle one strategy. **b**, shufflings based on *C*.*owc, H*.*cal, E*.*mue and R. esc* ortholog groups. In the blue bars, only *C*.*owc* ortholog locations have been shuffled. In the red bar series, all species ortholog locations were shuffled.

### 2.2 The outgroup comparison is the hard to pass test

The example with closely related genomes above suggests that the strategy of shuffling all gene to chromosome assignments has undesirable behaviour, reducing the data to totally independent observations even when there is underlying evolutionary structure.

Given the pairwise Oxford dot plots of ctenophore, sponge and cnidarian synteny, it is apparent that the gene locations of representatives of these species are highly correlated [14], and so likely affected by this problem. Further inspection of dotplots and the statistical significance of pairwise comparisons [14, 15] suggests that the critical test for whether data can be used to address the phylogenetic position of sponges or ctenophores is that of the significance of metazoan/unicellular outgroup combinations (compare SSR23 Extended Data Figs. 2 and 7). One way of looking at this problem is to consider each of the metazoan chromosome groups linked by shared orthologs as a hypothesis of an ancestral metazoan chromosome. We need not worry about the validity of the metazoan grouping in itself at this stage, as if it is not real it will not share orthologs with unicellular chromosomes in anything other than a random manner. Each of the observed metazoan groupings can be tested as a unit against the chromosomes of a unicellular genome, to determine if there is a significant enrichment of shared genes — if we randomly scattered all the unicellular orthologs of the metazoan grouping across the unicellular species chromosomes, how likely would we be to see the number we do see, or a larger number, by chance on our chromosome of interest? This probability under a null model (no significant enrichment) can be determined using the hypergeometric distribution (effectively Fisher’s exact test). For example in SSR23, the ctenophore/sponge/cnidarian chromosome group HCA7, EMU19, RES2 shares 15 orthologs with *Capsaspora* COW3. We can sum the total number of orthologs shared by the HCA7, EMU19, RES2 combination, with any *Capsaspora* chromosome, giving 27, and the number of orthologs on COW3 as 182. We also know the total number of orthologs in our data ^2^. Accordingly the uncorrected P-value for this comparison i.e. the probability of observing 15 or more orthologs on these chromosomes is, from the survival function of the hypergeometric distribution, 2.3e-09. Using this approach, we can calculate bespoke P-values for each hypothesis of metazoan-unicellular chromosome orthology without relying on a general threshold that does not consider the overall number of orthologs on a chromosome (which serves as a proxy for size). A multiple testing correction can then be applied, as each combination of chromosomes counts as a hypothesis tested. These values are compared to those in SSR23 in Table 1.

**Table 1:**
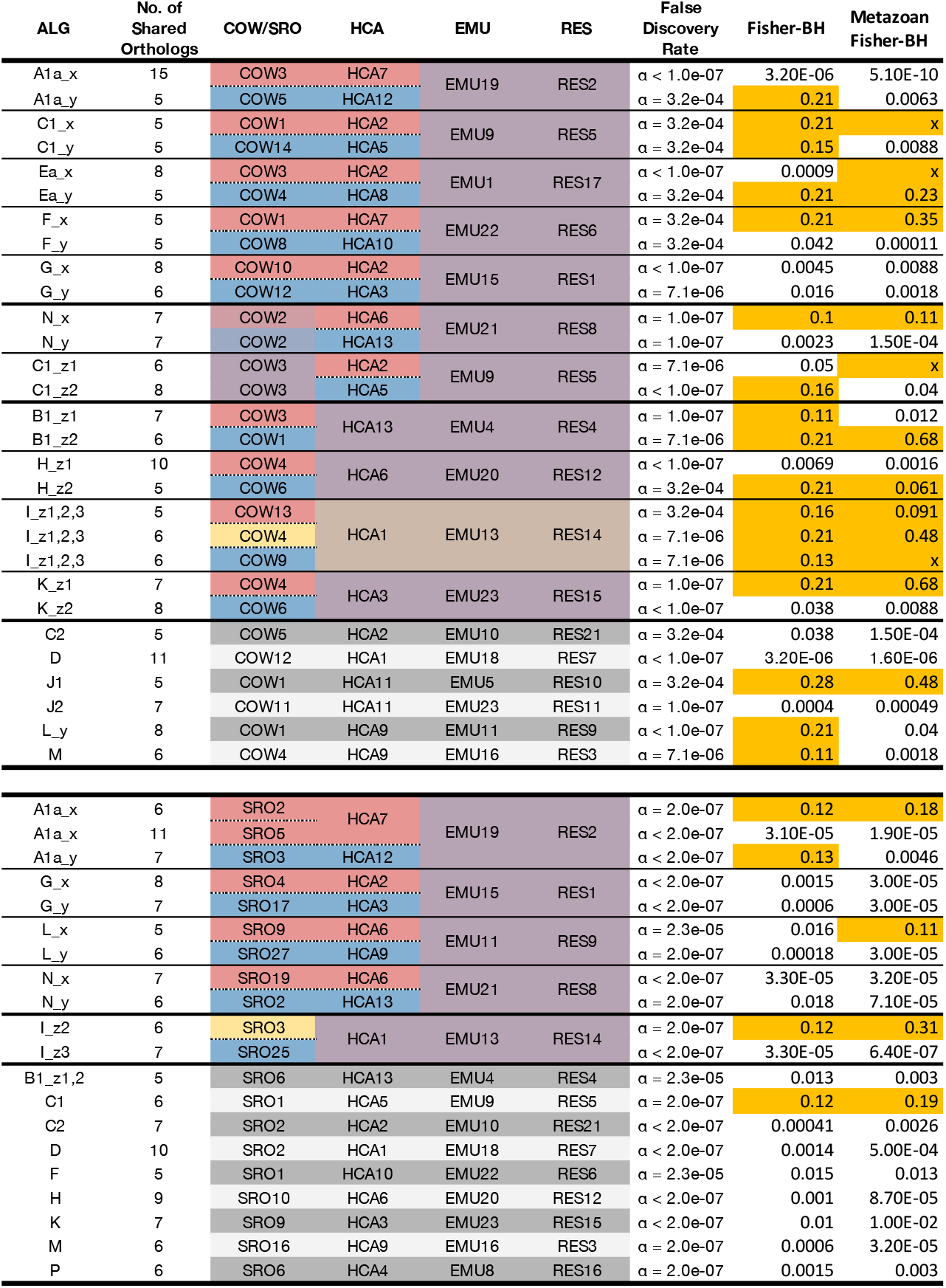
Recalculated significance of linkage groups for *S.ros* and *C.owc* with *H.cal, E.mue and R.esc* modified from SSR23. The ‘False Discovery Rate’ gives the SSR23 statistics. The column ‘Fisher-BH’ uses the orthology groups of SSR23 and gives FDRs calculated from Fisher P-values using the procedure described in the text and the Benjamini-Hochberg correction. The column ‘Metazoan Fisher-BH’ uses recalculated orthology groups and first identifies likely ancestral metazoan chromosomal combinations to reduce the number of tests, as described in the text. Where an ‘x’ is placed in this column, the metazoan chromosomal combination was not supported. Those highlighted in orange are no longer nominally significant (*α >* 0.05).

**Figure 2:**
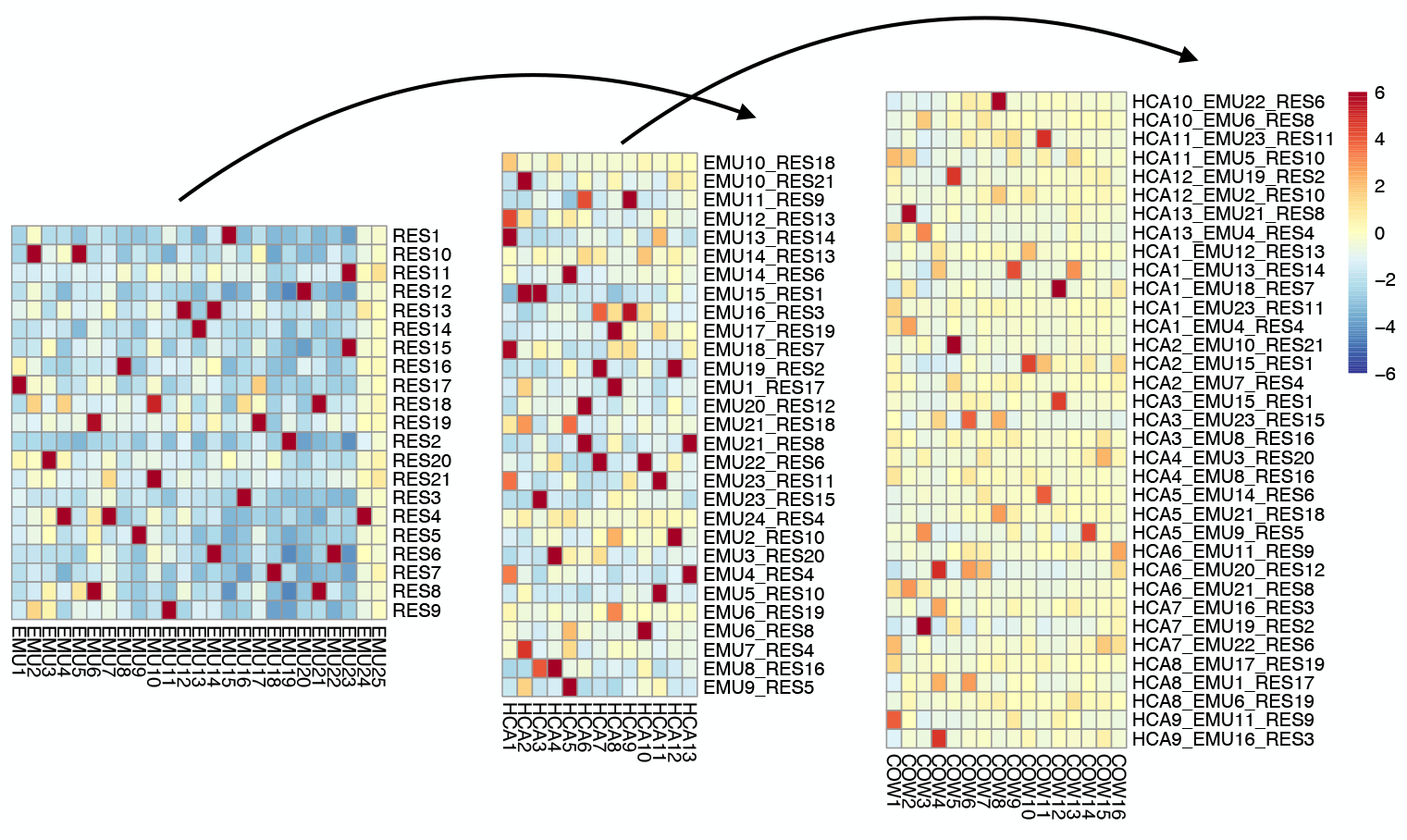
Example of the hierarchical build up of unicellular-metazoan syntenies. Each grid shows a heatmap representing counts of shared orthologs for all vs all chromosome (or combinations of chromosome) comparison, with all individual cells colored, on the same scale, by the standardized Pearson residuals for the omnibus chi-squared test represented in the heatmap. Significant syntenies between *Rhopilema* and *Ephydatia* (left) (the more red squares) are used as the basis for tests against *Hormiphora* (middle). The resulting syntenies are then compared to *Capsaspora* (right). Note that the distinction between true positives and noise becomes less clear across the comparisons, which is likely to lead to more acceptance of false positives, and/or rejection of true positives.

We can reduce the number of statistical tests that need to be corrected for by determining which metazoan chromosome combinations are likely to harbor ALGs, so that we only compare these composites to the chromosomes of the unicellular outgroup. I implemented a procedure to build up possible ancestral chromosomes for which there was strong statistical support, comparing first *Rhopilema* (cnidarian) and *Ephydatia* (sponge), then the results of that analysis with *Hormiphora* (ctenophore). These chromosomal combinations were then compared to the outgroup. This lets us make use of the greater number of orthologs found in the pairwise comparison between *Ephydatia* and *Rhopilema* and then within the metazoans when there is no unicellular outgroup, and so should *a priori* be favoured (Figure 2). The results are shown in the column ‘Metazoan Fisher-BH’ in Table 1.

### 2.3 The minor agreements

In *Capsaspora*, the genes XP_004365359.1 and XP_004365360.2 are at adjacent chromosomal locations and orthologs are found as adjacent pairs in *Ephydatia, Rhopilema* and Bilateria. They are not adjacent in *Hormiphora* but are on the same chromosome, giving a combination of COW1, HCA1, EMU20 and RES11. A similar situation is found with XP_004344437.1 and XP_004344438.2 or- thologs on COW12, HCA8, EMU18 and RES7. These combinations of chromosomes are not reported in SSR23 or detected as significant by the analyses above, but we can be sure that the gene pairs represent ancestral linkage groups because of the extreme unlikelihood of the adjacent gene pairing being found in all species for any reason other than by descent. If convergence driven by function is conjectured, then it is unclear why that should not also be possible with the examples of SSR23. Whereas it might be argued that the case of moving a small number of genes is not the same as a potential chromosome fusion or fission, that is also true for the ALGs of SSR23 which are based on numbers that are not dissimilar relative to the numbers of genes on a chromosome.

## 3 Discussion

I have argued here that SSR23 has overestimated the significance of the linkage groups that they have identified. Whether or not they are nominally significant, there is no way of experimentally testing whether syntenies supported by small numbers of genes are syntenic by descent or convergence. The re-analysis presented here shows ‘ALGs’ that, as far as comparisons with the unicellular outgroups go, look, at least in part, as if they were drawn from the right tail of a random probability distribution. Given this blurring of real syntenies and noise, it must be further remembered that absence of statistically supported evidence cannot be used to argue that further ALGs sharing few genes do not exist. Rather, from an examination of conserved adjacent pairs of genes, it seems that they do. We do not know how much these might affect the arguments of SSR23.

P-values and False Discovery Rates are designed to control type I error rates (false positives), but typically at the expense of higher type II errors (false negatives). This makes sense for e.g. GWAS studies with likely experimental follow up, but it is not obvious that phylogenetic signal should be preferentially concentrated in the small number of linkages supported by more robust synteny groups. It is reasonable to conclude from Figure 2 that the overlap between signal and noise increases with phylogenetic distance.

Given this, the use of syntenies that are supported by few genes in phylogenetic debate should be viewed critically, in line with any other sort of evidence.

## 4 Methods

### 4.1 Sequence data

For *Capsaspora, Salpingoeca, Hormiphora, Ephydatia* and *Rhopilema*, protein sequence predictions and chromosomal locations were taken from the supplementary information of SSR23. The A form of *Capsaspora* was used, noting SSR23’s observation that this choice makes no difference to results. Gene order and protein sequences for the homoscleromorph sponge *Corticium candelabrum* and the anthozoan cnidarian *Nematostella vectensis* were taken from the NCBI (accessions GCF_963422355.1 and GCF_932526225.1).

### 4.2 Ortholog identification

I calculated 4-way orthologs using the `ssearch3` program, an implementation of the Smith-Waterman algorithm from the FASTA package [16]. Reciprocal best hits, with an E-value < 0.1 for pairwise comparisons were collected and cross-referenced to produce fully consistent 4-way species clusters i.e. for each ortholog group, single genes from all 4 species were each others best hits (see github code repository for implementation details). Where stated I used the 4-way orthologs provided in the supplementary information of SSR23: `COW_EMU_HCA_RESLi_reciprocal_best_hits.rbh` and `EMU_HCA_RESLi_SRO_reciprocal_best_hits.rbh`. Both approaches produce similar results.

### 4.3 Chromosome comparisons

For all 4-way orthologs, an outgroup (the single celled eukaryote, so either *Capsaspora* or *Salpingoeca*) and an ingroup, namely the metazoans *Hormiphora, Ephydatia* and *Rhopilema* were defined. For each orthologous group, chromosome ids were retrieved for the 4 genes. A table of of ingroup and outgroup chromosome counts was created, where the 3 ingroup chromosome ids were combined to give one identifier (e.g. the three ids HCA7 RES2 EMU19 became the single id HCA7_RES2_EMU19), iterating over all orthologous groups. Then, for each unique combination of 4 chromosomes with shared orthologs (e.g. COW3, with HCA7_RES2_EMU19, *N* orthologs), the total number of observations of that combination, the total number of orthologs on COW3, the total number of orthologs on HCA7_RES2_EMU19 combinations and the total number of 4-way orthologs in our dataset were retrieved. An exact probability of seeing *N* or more observations was calculated using the hypergeometric distribution. The set of P-values for all pairwise comparisons was then corrected for multiple testing using the Benjamini-Hochberg procedure.

This procedure was generalized to allow 4-way comparisons to be built up from 2-way and 3-way comparisons, restricting 4-way comparisons to chromosome combinations that had been judged significant in 3-way comparisons, and 3-way comparisons to those significant in 2-way, as illustrated in Figure 2. The standardized residuals used in this figure were calculated using the `standardized_resids` method of the python `statsmodels` module, but are not used in the main calculations.

Code and data are available at https://github.com/rcply/synteny

### 4.4 Conserved adjacent pairs of orthologs

Conserved adjacent pairs of orthologs were identified using the approach described in [13].

## 5 Acknowledgements

Many thanks to Iain Mathieson (University of Pennsylvania), Max Telford (UCL), Graham Budd (Uppsala), Davide Pisani (Bristol), Evelyn Houliston and Stefania Castagnetti (Villefranche-sur-mer) for helpful discussions.

## Footnotes

1 That within all species, all gene locations across all chromosomes are shuffled in SSR23 permutation tests is not completely obvious from the method text, which refers to “10 million permutations of gene indices” (P.58 supplementary information [14]), but it is apparent from the archived code `odp_nway_rbh` (https://zenodo.org/records/7857390 lines 415-6), where each `thisscaf` is a dataframe column representing a species.

2 Giving ((15, 167), (12, 1680)) as a contingency table.

